# Specific determinants of the Transmembrane region of the Andes virus Gc glycoprotein drive the transition from membrane hemifusion to pore formation

**DOI:** 10.64898/2026.03.17.712151

**Authors:** Chantal L. Márquez, Fernando Villalón-Letelier, Gianina Arata-Salas, Nicole D. Tischler

## Abstract

Andes virus (ANDV), a highly pathogenic orthohantavirus, enters host cells through low pH–triggered membrane fusion mediated by the Gc glycoprotein, a class II fusion protein containing a single C-terminal transmembrane domain (TMD). While the ectodomain has been extensively characterized, the role of the TMD in late-stage fusion remains unclear. Here, we investigated the minimal functional length and sequence requirements of the ANDV Gc TMD using site-directed mutagenesis. C-terminal deletion mutants and serine-to-alanine substitutions were evaluated for protein expression, virus-like particle production, cell–cell fusion, pseudotyped vector entry, and hemifusion activity. Deletion of the Gc cytoplasmic tail (CT) or a single C-terminal TMD residue was tolerated, whereas deletion of two or more residues impaired particle production and fusion, indicating that at least 21 of the 22 TMD residues are required for efficient membrane fusion and viral entry. Hemifusion assays showed that deletion of two or three residues, or substitution of the strictly conserved S1121, allowed lipid mixing but blocked progression to full fusion, while deletion of four residues also abolished hemifusion. In contrast, mutation of the less conserved S1126 had minimal effect. These results identify a precise TMD length and a conserved polar TMD residue as critical determinants of fusion pore formation in ANDV.

## INTRODUCTION

Orthohantaviruses are enveloped, tri-segmented RNA viruses classified as a separate genus within the Hantaviridae family (Bunyaviricetes class) [1]. This group includes several human pathogenic viruses with worldwide distribution and persist in small mammals, primarily in rodents, by establishing persistent infections. Their transmission to humans occurs through inhalation of aerosolized, contaminated excreta [2,3]. Andes virus (ANDV), endemic in Chile and Argentina, is unique among orthohantaviruses in that human-to-human transmission has been documented, and hence represents an enhanced threat to public health [4,5]. Infections with ANDV can result in hantavirus cardiopulmonary syndrome (HCPS) with fatality rates up to 40% [6].

Orthohantaviruses enter into the host cells through receptor binding and uptake into endosomes, where viral and endosomal membrane fusion must occur to allow the virus access to the cytoplasm where the replication of the virus takes place [7]. This process is driven by the viral Gn and Gc glycoproteins organized into tetrameric spikes on the viral surface. While Gn is likely involved in receptor binding, Gc drives the membrane fusion process activated at low pH within endosomes [8–10]. Gc shares structural features with class II fusion proteins composed of three β-rich domains (DI, DII and DIII), a membrane proximal region (stem region), a C-terminal transmembrane domain (TMD) and a short cytoplasmic tail (CT) [11,12]. The membrane fusion process involves multiple consecutive steps, which are the exposure and anchoring of the target membrane insertion surface (TMIS) into the endosomal membrane, apposition of the viral and host membranes, merger of the outer membrane monolayers (referred to as hemifusion), lipid mixing of the inner membranes to form the fusion pore and finally the extension of the membranous pore [13].

While the conformational changes of the ectodomains of class II fusion proteins have been extensively characterized, the contribution and requirements of the TMD in late-stage fusion events remains largely underexplored. A significant number of studies based on the TMD of class I and class III fusion proteins have shown that this region is not merely a membrane anchor, but it fulfills crucial roles during membrane fusion, providing structural flexibility during conformational translations, promoting TMD-TMD oligomerization and fusion protein trimerization, interacting with the TMIS and inducing lipid mixing and fusion pore formation (nicely reviewed in [14–16]). In class II-enveloped viruses such as those of flavi-alpha- and pestiviruses, much less information is available. While for alphaviruses the interactions between the TMDs of the E1 fusion protein with the E2 companion protein seems to be essential for viral assembly and membrane fusion during cell entry [17,18], it was not possible to determine a specific sequence requirement [19]. Flaviviruses in turn, contain two antiparallel TMD helices that form a coiled-coil, that are further coordinated with the paired TMDs of the companion protein, forming membrane-embedded complexes that regulate the prefusion stability of the spikes [20–23]. Structural studies of the TMDs *in situ* embedded in the viral membranes of dengue and Venezuelan equine encephalitis viruses have shown that glycine or serine residues are located at TMD interfaces, while hydrophobic residues face outwards [20,22]. A mutagenesis study of the flavivirus E fusion protein further demonstrated the requirement of the two antiparallel TMDs to induce late membrane fusion steps and homotrimer stabilization [21]. Different to class I fusion proteins, a minimal TMD length to support membrane fusion has not been defined for class II fusion proteins.

In contrast to flavi- and alphaviruses, the Gc protein of orthohantaviruses and other class II-enveloped bunyaviruses contains a single TMD, whereas their companion protein Gn is anchored by two transmembrane helices [24]. It remains unclear whether the single Gc TMD includes specific features to support membrane fusion. In this study we aimed to determine the minimal functional length of the ANDV Gc TMD and to test for the role of conserved, polar residues within this region. We found that deletion of the Gc-CT and one residue from the C-terminus of the TMD was tolerated, allowing full membrane fusion, whereas deletion of 2 and 3 residues arrests fusion at the hemifusion stage. In parallel, substitution of a strictly conserved serine residue also blocks progression to full fusion without abolishing outer leaflet lipid mixing. These findings demonstrate the precise TMD length and a single conserved polar residue are critical determinants of the transition from hemi-fusion to fusion pore formation. Our results extend current models of class II fusion by revealing that late-stage fusion is governed by sequence features within the single Gc transmembrane helix.

## MATERIALS AND METHODS

### Cells and antibodies

Vero E6 cells (ATCC, CRL 1586) and 293FT cells (Invitrogen) were maintained in Dulbecco’s modified Eagle’s medium supplemented with 10% FBS (Gibco). CHO-K1 cells were maintained in Ham’s F-12K (Kaighn’s) medium supplemented with 10% FBS (Gibco). Monoclonal antibodies (mAbs) against ANDV antigens were developed previously in our laboratory and included anti-Gn 6B9/F5 [25] and anti-Gc 2H4/F6 [26]. The anti-Gc mAb 5D11/G7, also generated in our laboratory, was used to corroborate all results.

### Mutagenesis

Several mutants within the glycoprotein precursor (GPC) of ANDV (Orthohantavirus andesense) were generated by DNA mutagenesis using Pfx polymerase (Invitrogen)-driven PCR amplification. The TMDs of Gn and Gc were predicted by TMHMM prediction server (http://www.cbs.dtu.dk/services/TMHMM-2.0/) [27], suggesting residues 1108-1129 for the TMD of Gc. Deletion mutants of the C-terminal residues of the TMD were generated by introducing translation termination codons into the ANDV GPC coding region (Gen bank accession number AY228238.1), hence also eliminating the Gc CT. The site-directed mutation of S1121A was generated by primer-introduced mutagenesis. All primers are listed in Table 1. Subsequently, the PCR products were subcloned into the pI.18 expression vector using BglII and XhoI restriction enzymes (Invitrogen) and positive clones corroborated by sequencing (Macrogen Inc.). The site-directed mutant S1126A was generated through a mutagenesis service (Genscript Inc.). The deletion mutant of the Gc CT (Gc-ΔCT) has been constructed and characterized previously [28].

**Table 1.**
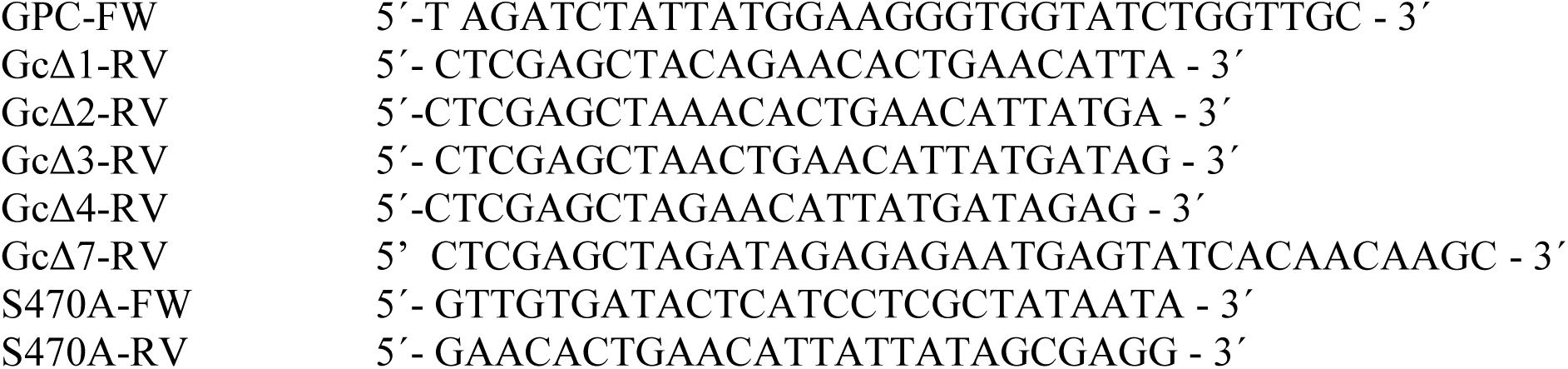
Primers used for the preparation of Gc TMD deletion and substitution mutants.

### Biotinylation

Detection of cell surface proteins on transfected cells was performed by cell surface biotinylation using a cell-surface protein isolation kit (Pierce). Briefly, 293FT cells in 100 mm plates were transfected with 8 μg of DNA using the calcium phosphate method [29], and 48 h post-transfection, the cell surface proteins were biotinylated following the manufacturer’s instructions. Gc proteins were detected in the biotinylated (surface proteins) and non-biotinylated (intracellular proteins) fractions by western blotting using mAb anti-Gc 2H4/F6 at a dilution of 1:2.500 and β-actin was detected with the mAb anti-β-actin (Sigma) at 1:5.000 as a control. Anti-mouse immunoglobulin HRP conjugate (Sigma) was used to detect the primary antibody at 1:5.000 and the detection of HRP was performed with a chemiluminescent substrate (SuperSignal WestPico; Pierce).

### Cell–cell fusion assay

This assay was performed as described previously [30]. Vero E6 cells seeded into 16-well chamber slides (NUNC) were transfected with 0.5 μg of plasmids DNA encoding GPC, including WT or mutant Gc, using lipofectamine 2000 (Invitrogen) as indicated by the manufacturer. 48 h post-transfection, fusion was triggered by incubating the cells with low pH medium (E-MEM pH 5.5) at 37 °C for 5 min. 3 h later, the cells were stained for 1 h with 1 μM of CellTracker CMFDA (Invitrogen), fixed with 4% paraformaldehyde for 20 min and permeabilized with 0.1% Triton X-100 in PBS for 15 min at room temperature. Finally, cells were incubated for 1.5 h with the mAb anti-Gc 5D11/G7 at 1:500, and next with secondary antibody anti-mouse immunoglobulins conjugated to Alexa555 (Molecular probes, Invitrogen) at a 1:500 dilution for 1 h followed by nuclei stain with DAPI 1 μg/ml in PBS for 5 minutes. Samples were examined under a fluorescence microscope (BMAX51; Olympus) and pictures taken for subsequent quantification (ProgRes C3; Jenoptics). For each sample, the fusion index was calculated using the formula: 1-[number of cells/number of nuclei] and the mean fusion index of at least three fields per experiment was calculated.

### Cell-based hemifusion assay

The cell-based hemifusion assay was performed as previously described [31] with some modifications. Effector GM1+ 293FT cells seeded into 16-well plates were transfected with 5 μg of plasmid using lipofectamine 2000 (Invitrogen) as indicated by the manufacturer and 45 h post-transfection were detached from the plates using diluted trypsin for 3 min. Effector cells were centrifuged at 700 x g for 5 min, resuspended in DMEM, and counted. At the same time, target GM1-CHO-K1 cells were trypsinized and subsequently labeled with the Celltracker blue CMAC (Molecular Probes) at 40 μM in F12-K medium for 40 min at 37°C. Cells were centrifuged at 700 x g for 5 min and incubated with fresh F12-K medium for 30 min in order to eliminate the excess of dye. After washing with PBS, target cells were resuspended in DMEM and counted. Effector and target cells were co-cultivated into 16-well chamber slides at ratio 1:4 for 3 h and next the fusion process was activated by incubation with a low pH medium for 5 min at 37°C (DMEM, pH 5.5). The medium was subsequently replaced with DMEM (pH 7,2) and 30 min later the cells were fixed with 4% PFA. For immunofluorescence staining, the samples were next incubated for 30 min with 5 μg/ml Alexa Fluor 488-conjugated cholera toxin B subunit (CTX) (Molecular Probes) at 37°C, washed with PBS and mounted with DABCO. Ten confocal images for each condition were collected using a confocal fluorescence microscope (Olympus Fluorview FV1000) and the percentage of GM1 transference was calculated using the formula: [number of (CMAC+-CTX+ cells / number of CTX+ cells that were in contact with at least one target cell].

#### Production of virus-like particles

VLPs were produced as described previously (Acuña et al., 2014). Briefly, pI.18 plasmids constructs encoding WT ANDV GPC or mutant GPC were used to transfect 293FT cells using the calcium phosphate method. 48 h post-transfection, cell supernatants were collected and clarified by sucrose sedimentation and resuspended in HEPES-NaCl-EDTA (HNE) buffer.

Production of pseudotyped simian immunodeficiency (SIV) vectors and transduction. SIV vectors pseudotyped with ANDV Gn and WT or mutant Gc were produced as described (Cifuentes-Muñoz et al., 2010). 293FT cells were co-transfected with pSIV3+ [79], pGAE1.0 [80], and pI.18/GPC encoding WT or mutant Gc. Supernatants were harvested 72 h post-transfection, concentrated by ultracentrifugation (135,000 x g), and used to transduce Vero E6 cells. After 72 h, GFP reporter gene expression was analyzed by flow cytometry (BD FACSCanto II, BD Biosciences). Normalized transduction was calculated from the percentage of GFP-positive cells (≥10,000 cells counted per condition).

### Statistical analysis

Data from at least three independent experiments are shown as mean ± standard error of the mean (SEM). Statistical significance was determined using an unpaired Student’s t-test in GraphPad Prism v6.0. For normalized samples, one sample Wilcoxon test was applied, using a theoretical median of 100. Significance thresholds were defined as follows: P < 0.05 (*), P < 0.01 (**), and P < 0.001 (***).

## RESULTS

### Design, synthesis and transport to the cell surface of Gc TMD mutant proteins

To determine the minimum length of the ANDV Gc TMD required to preserve fusion activity, a panel of mutants was designed. Therefore, we analyzed the recently published *in situ* structures of the ANDV Gn/Gc spikes and their TMDs by Guo et al. [32]. The Gc TMD forms a straight helix which is positioned adjacent to Gn-TMD2, establishing hydrophobic contacts between both helices (Figure 1) [32].

**Fig 1.**
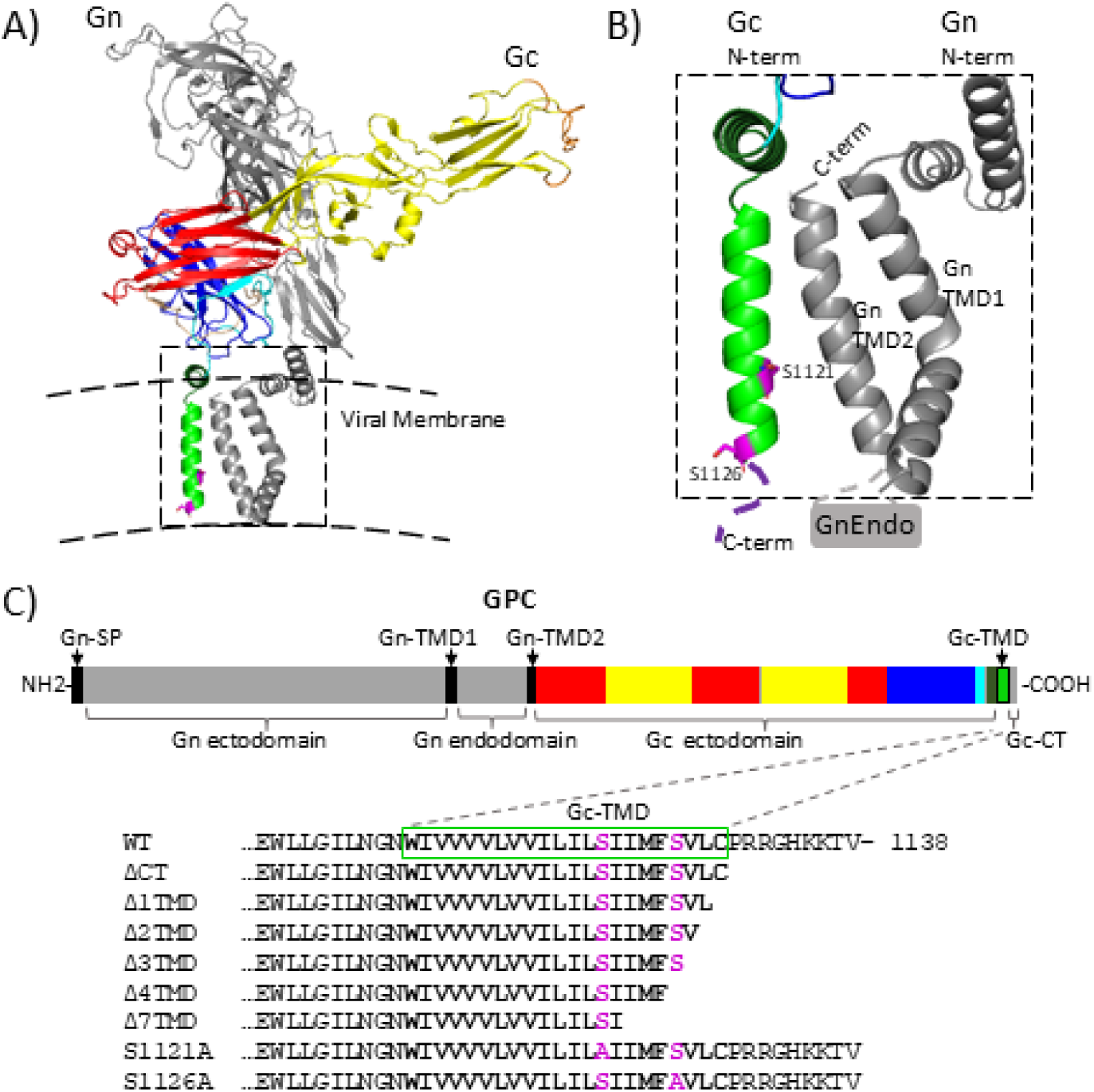
High-resolution in situ ANDV Gn/Gc structure embedded in virus-like particles in cartoon representation and schematic representation of Gc TMD mutants within the glycoprotein precursor (GPC). (A) Structural model of the cryo-EM electron density map of the Gn/Gc heterodimer obtained from Andes virus-like particles shown as ribbon representation (EMDB: 9P3X) [32]. The domains are color-coded as follows: Gn ecto- and endodomain, grey; Gc domain I, red; Gc domain II, yellow; Gc domain III, blue; Gc stem region, cyan; Gc pre-transmembrane domain (pre-TMD) region, dark green; Gc transmembrane domain (TMD), neon green. The viral membrane is represented by dashed black lines. (B) Enlarged view of the pre-TMD and TMDs of Gn and Gc, shown in the same color scheme as in (A). Residues subjected to site-directed mutagenesis are highlighted in magenta. The Gc cytoplasmic tail (CT) is schematized in dark purple. The N- and C-termini of Gn and Gc are indicated. The Gn endodomain (GnEndo), which was not resolved in the structure, is depicted schematically as a boxed region. (C) Schematic organization of the glycoprotein precursor (GPC), using the same color scheme as in (A). The N-terminal signal peptide (SP) and the TMD1 and TMD2 regions of Gn are indicated in black. The C-terminal sequence of Gc WT is shown, including the TMD (outlined in a neon green box) and the CT. Residues targeted for site-directed mutagenesis are highlighted in magenta. Below the WT sequence and the corresponding C-terminal sequences of each mutant construct are displayed.

In addition, we examined residue conservation within the Gc TMD of the genus Orthohantavirus and the entire Hantaviridae family. Within orthohantaviruses, only two polar residues are present in the TMD; S1121 and S1126. Both residues are located within the C-terminal half of the TMD and are therefore predicted to be located within the inner leaflet of the viral membrane (Figure 1). The residue S1121 is strikingly conserved within the Orthohantavirus genus and across all other genera of the Hantaviridae family, with the exception of the two genera including species that persist in piscine hosts. In contrast, S1126 is only partially conserved within the Orthohantavirus genus, where conserved mutations to threonine are observed, and is additionally conserved in three other genera within the family (Figure S1). Based on these structural and conservation analyses, we introduce C-terminal deletions in the Gc TMD as well as alanine substitutions of the conserved polar serine residues. Specifically, we generated mutants lacking 1, 2, 3, 4 or 7 C-terminal residues of the Gc TMD C-terminal, including the Gc CT, and constructed the single- residue substitutions mutants S1121A and S1126A (Figure 1C).

To evaluate if these Gc deletion mutants were properly synthesized and folded, we assessed their expression and accumulation at the cell surface. To this end, 293FT cells were transfected with plasmids encoding GPC WT or the respective mutants. At 48 h post-transfection, the cell supernatant was collected to assess the presence of virus-like particles (VLPs), and the cells were subjected to cell surface protein biotinylation. After concentration of the cell supernatants and separation of the intracellular and biotinylated fractions, all samples were analyzed by western blot (Figure 2). β-actin immunodetection served as loading control in the intracellular fraction, and as a negative control of the biotinylated fraction, confirming cellular integrity. We detected Gc WT and all mutants with anti-Gc mAb in the intracellular and surface fractions. In the intracellular fraction, which were prepared for higher electrophoretic separation, the TMD deletion mutants displayed a shift in their electrophoretic mobility compared with Gc WT (55 kDa), consistent with the respective amino acid deletions of 10 (Δ1TMD), 11 (Δ2TMD), 12 (Δ3TMD), 13 (Δ4TMD) and 16 (Δ7TMD) residues. All mutants were present in intracellular and surface fractions in levels comparable to WT; although Δ7TMD showed lower amounts in one biological replicate, the quantification of several biological replicates showed no significant difference between Gc WT and any of the TMD mutants in both fractions (Figure 2B).

**Fig 2.**
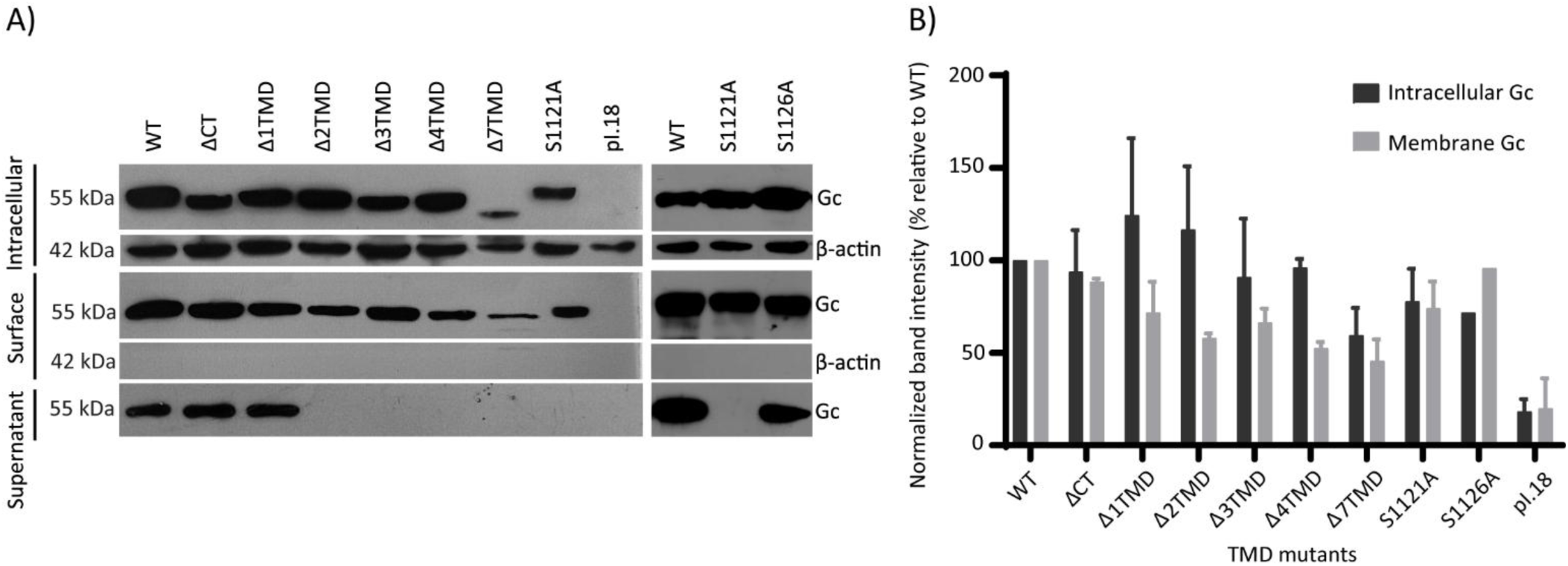
Expression, cell surface localization and VLP release of ANDV GPC including Gc TMD deletions or mutants. The supernatant of 293FT cells expressing WT or mutant GPC was harvested and concentrated 48 h post-transfection, and cells surface-biotinylated. Subsequently, intracellular and surface protein fractions were prepared and all samples subjected to Western blot analysis with mAb anti-Gc (top) and β-actin (bottom). Quantification from N = 2 to 4 experiments, depending on the mutant. Statistical analyses were performed using one sample Wilcoxon test. No significant differences between WT and Gc mutants were observed.

β-actin was only detected in the intracellular fractions of all samples, confirming the preservation of cellular integrity during the biotinylation process. Analysis of the concentrated cell supernatant revealed detectable Gc only for WT, Δ1TMD and S1126A, whereas no signal was observed for the remaining mutants. Because the detection of Gc in the cell culture media correlates with VLP production [11,28,33], these results suggest that deletion of more than 1 residue from the TMD, as well as the TMD mutation of residue S1121, impairs virus particle assembly and/or release.

### Cell-cell fusion and cell entry activity of Gc TMD mutants

We next analyzed the fusion activity of the TMD mutants, using a cell-cell fusion assay [30]. We also included the GPC mutant ΔCT for comparison, for which we have previously shown cell surface accumulation similar to WT levels [28]. Vero E6 cells were transfected with plasmids encoding either WT or mutant GPC. After 48 h post-transfection, cells were exposed during 5 min to low pH to trigger activation of the Gc fusion protein. Cells expressing WT Gc and exposed to low pH exhibited numerous syncytia containing approximately 4 to 20 nuclei (Figure 3A, white arrowheads).

**Fig 3.**
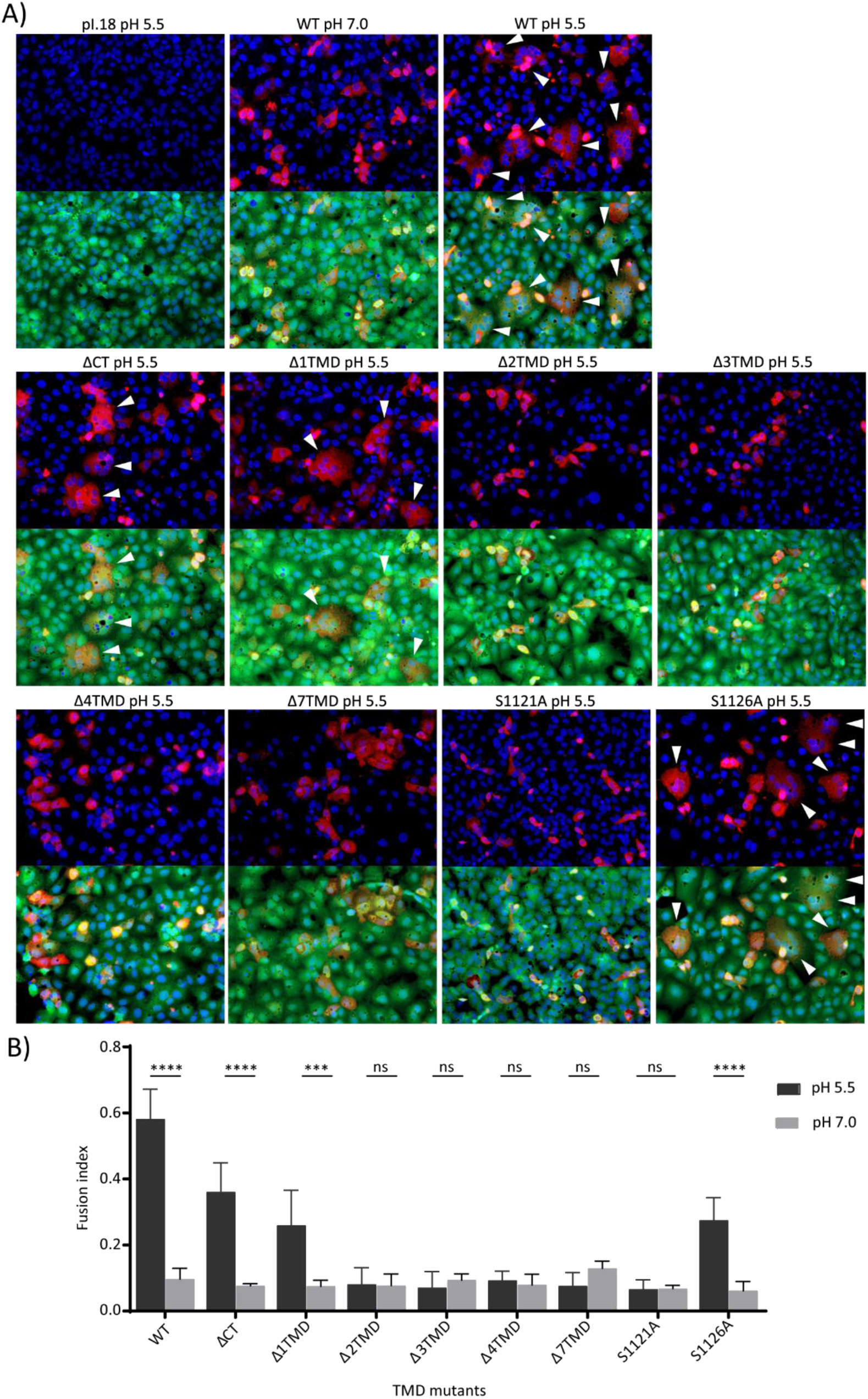
Cell-cell fusion activity of ANDV GPC WT and mutants and quantification. A) Vero E6 cells expressing WT or mutant Gc were exposed to low pH and analyzed for syncytia formation by fluorescence microscopy with a 20x objective magnification. Cytoplasm, CMFDA (green); Nuclei, DAPI (blue); Gc, Alexa-555 (red). Results representative of at least N=3 biological repeats. B) Cells and nuclei were counted in at least five microscope fields per sample for each condition and the fusion index calculated. Quantification from at least N=3 independent experiments. Statistical analyses were performed using an unpaired Student’s t-test; P < 0.05 (*), P < 0.01 (**); P < 0.001 (***); ns, not significant.

These multinucleated structures were not observed in cells maintained at neutral pH (Figure 3A). Among the mutants tested, only ΔCT and Δ1TMD promoted syncytium formation following low-pH treatment. However, in both cases, syncytia were less frequent and generally smaller than those observed in WT Gc-expressing cells (Figure 3A). These results are reflected in the quantitative analysis shown in Figure 3B. Quantification revealed a fusion index of around 0.6 was observed for WT under low-pH conditions (dark grey bars), whereas ΔCT, Δ1TMD and S1126A exhibited a reduced fusion index ranging between ∼0.25-0.4. In contrast, mutants including more than two residue deletions of TMD and point mutations S1121A displayed fusion indices without statistical difference compared to samples maintained at neutral pH (light gray bars), indicating a complete loss of fusogenic activity. Together, these findings indicate that the Gc-CT is not strictly required for membrane fusion but contributes to optimal fusogenic activity. In contrast, the Gc TMD is essential for fusion, as at least 21 of the 22 residues are required to preserve activity. Furthermore, the substitution of the less conserved, and more C-terminally located Ser1126 retained fusion activity, while the strictly conserved Ser1121 residue plays a crucial role in membrane fusion.

We corroborated these findings by introducing the same mutations onto SIV–based pseudotype vectors bearing the ANDV Gn/Gc spikes. In this previously developed system [25], viral entry can be monitored through expression of a fluorescent reporter gene. Prior to transduction assays, we analyzed concentrated supernatants from 293FT producer cells by western blot to assess incorporation of Gn/Gc onto the pseudotyped vectors. Consistent with the VLP assembly results, pseudotyped vectors were detected only for ΔCT, Δ1TMD and S1126A (Figure 4A). The remaining mutants were not detectable in the supernatant, despite comparable levels of the SIV p28 capsid protein in all conditions, indicating that SIV particle production per se was not impaired. Subsequent transduction of Vero E6 cells with these pseudotyped viral vectors revealed that all three detectable mutants exhibited reduced entry efficiency, with transduction levels reaching only ∼25–40% of WT (Figure 4B, Figure S2). These results closely mirror the reduced fusion activity observed in the cell–cell fusion assays (Figure 3).

**Fig 4.**
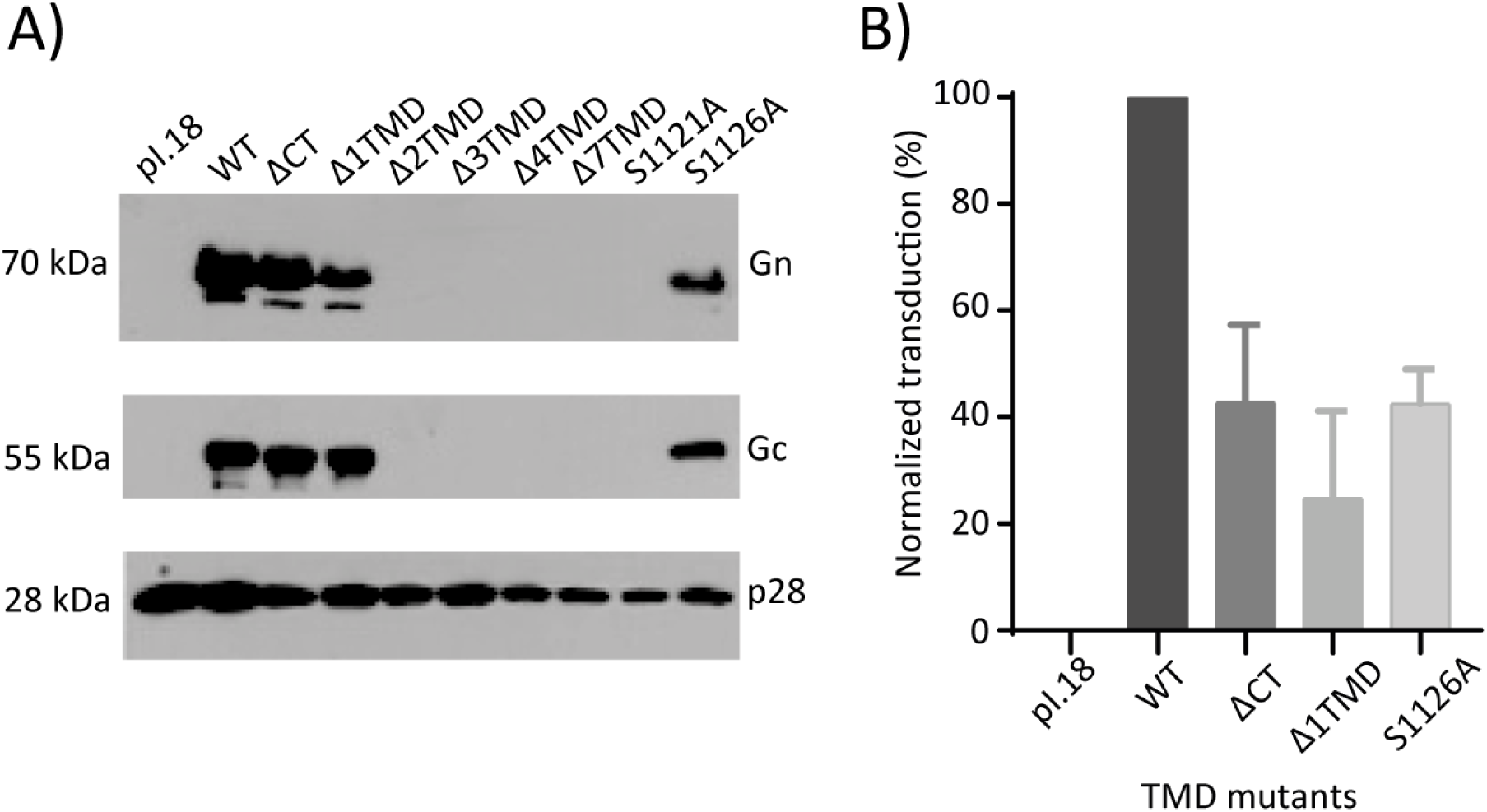
Preparation of SIV vectors pseudotyped with ANDV WT or Gc TMD mutants and quantification of cell transduction. A) Western blot of the concentrated cell supernatant of 293FT transfected with the different plasmids 72 h post-transfection using anti-Gn, anti-Gc and anti-SIV p28 mAbs. B) Normalized mean values of transduced Vero E6 cells with SIV vectors pseudotyped with empty pI.18 vector or ANDV GPC WT or Gc TMD mutants. Experiments are representative of N = 2 replicates.

### Hemifusion activity of Gc TMD mutants

The cell–cell fusion assay measures syncytium formation, a process that requires complete membrane merger, including fusion pore formation and expansion. To obtain molecular insights into which specific steps may be blocked in fusion inactive mutants, particularly late fusion steps, we established a cell-based hemifusion assay adapted from Giraudo et al. (2005), which detects lipid mixing between the outer leaflets of adjacent plasma membranes. Hemifusion was monitored by measuring transfer of the ganglioside GM1 from 293FT effector cells to CHO-K1 target cells. 293FT cells accumulate GM1 in the outer leaflet of the plasma membrane [34] whereas CHO-K1 cells lack GM1 due to a deficiency in its biosynthetic pathway [35]. Effector cells transfected with ANDV WT or mutant GPC were co-cultured with CMAC-labeled CHO-K1 target cells. Following low-pH treatment, GM1 transfer was detected using Alexa Fluor 488–conjugated cholera toxin B subunit (CTX), which specifically binds GM1 [36].

As shown in Figure 5A, the Gc-Δ2TMD, Gc-Δ3TMD and S1121A mutants promoted GM1 transfer from effector (green) to target cells (blue), indicating that these mutants retain the ability to induce lipid mixing. However, not all effector cells mediated transfer, likely reflecting through the transfection efficiency which was on average 32.4%. In contrast, for the Gc-Δ4TMD mutant, membrane transfer was not apparent. Quantification of GM1 transfer under low-pH conditions revealed levels of approximately 70% for WT, Gc-Δ2TMD, Gc-Δ3TMD and S1121A (Figure 5B, dark gray bars). In comparison, the transfection of the empty vector led to a basal transfer level of ∼20%, which was observed in all neutral pH conditions, reflecting the non-specific ganglioside exchange due to high cell density and handling procedures. Importantly, the increase in lipid transfer observed for Δ2TMD, Δ3TMD and S1121A compared with neutral pH controls was statistically significant. In contrast, Δ4TMD exhibited under low pH activation ∼40% GM1 transfer with a high standard deviation between experiments, lacking statistical significance with its neutral pH control (Figure 5B).

**Fig 5.**
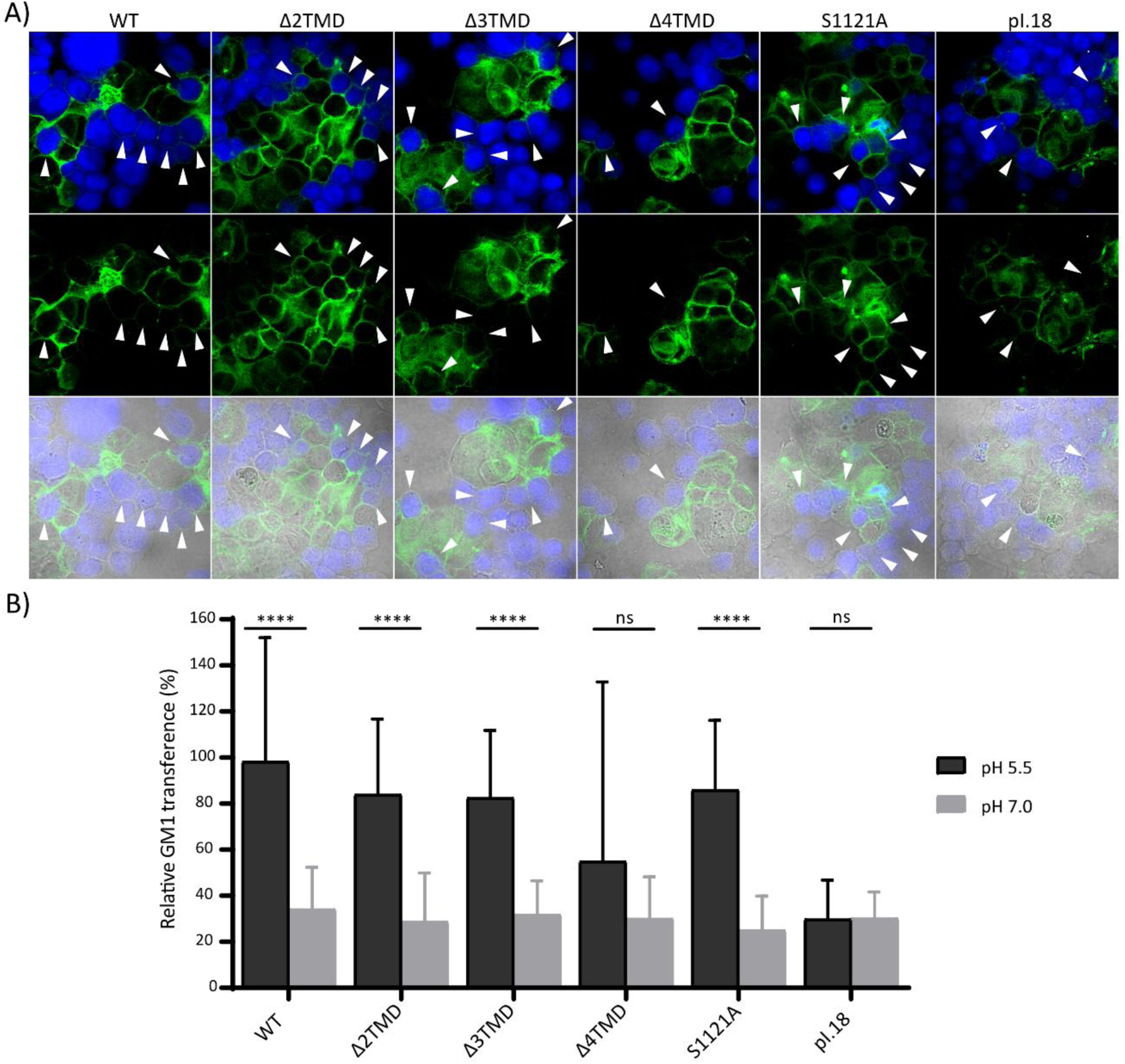
Cell-based hemifusion assay and quantification. A) Confocal microscopy 293FT cells stained with the membrane dye CTX-Alexa 488 (GM1+, green) (effector cells) co-incubated with CMAC-labeled CHO-K1 cells (target cells, blue). Images were acquired using a 40x magnification objective. White arrowheads indicate membrane dye transfer to CMAC-labeled CHO-K1 cells. B) Quantification of the hemifusion activity by counting cells that present both dyes (GM1+/CMAC+). The hemifusion percentage was determined using the formula [100*(GM1+/CMAC+)/(GM1+)]. Experiments are representative of N = 4 independent replicates with ten microscopy fields acquired per condition of each experiment. Statistical analyses were performed using an unpaired Student’s t-test; P < 0.05 (*), P < 0.01 (**), P < 0.001 (***), P<0.0001 (****); ns, not significant.

Collectively, these findings demonstrate that progressive shortening of the Gc TMD differentially affects distinct stages of membrane fusion. Deletion of two or three residues permits lipid mixing and hemifusion but prevents efficient fusion pore formation or expansion, whereas deletion of four residues completely abrogates hemifusion. Furthermore, the polar residue S1121 also plays a crucial role in the late fusion step stage, promoting the transition from hemifusion to full fusion. These data highlight a critical and length-dependent role of the ANDV Gc TMD domain in driving the membrane fusion process.

## DISCUSION

Membrane fusion is a critical step in the viral replication cycle, enabling delivery of the viral genome into the host cell cytosol. Given that specific features of the TMD influence the activity of several viral fusion proteins, we examined the role of the single TMD of orthohantavirus Gc in the virus particle assembly and membrane fusion, using ANDV as a model. Our analysis reveals strict requirements for both, TMD length and a specific serine residue.

Truncation experiments of the Gc TMD showed that mutants spanning at least 21 of the 22 residues still allowed syncytia formation and pseudotyped virus cell entry, although with reduced efficiency, similar to mutants with the deletion of the Gc-CT alone. Deletion of a single residue reduces but does not abolish fusion activity, suggesting that both, the Gc-CT and subtle changes in the Gc TMD length can influence fusion efficiency. The C-terminal residue of the Δ1TMD mutant corresponds to a cysteine residue that could potentially undergo lipid modification such as palmitoylation, a modification known to occur in influenza hemagglutinin (HA) and to influence fusion efficiency [37,38]. Although speculative, the partial reduction in fusion activity observed with both mutants may reflect altered membrane interactions or positioning of the TMD.

Deletion of two or three Gc TMD residues arrested the mutants in a hemifusion intermediate state, where lipid mixing occurred without pore formation, while deletion of four residues abolished significant lipid mixing activity. Minimal length requirements have been reported for several class I fusion proteins of viruses; 18 of 25 residues for human immunodeficiency virus (HIV) gp41 [39–41], 17 of 27 residues for influenza virus HA [42], and 23 of 26 residues for SIV gp41 [42,43]. The length requirement observed for ANDV Gc therefore appears more stringent, suggesting that a precise TMD length is particularly important for orthohantavirus fusion proteins.

The hemifusion phenotype of the ANDV Gc TMD mutants was established using the cytoplasmic dye CMAC of Mr= 210 KDa, which can penetrate small pores. Mutants lacking two or three residues resemble truncated HA constructs, in which deletion of 12 residues from either end of the TMD allowed lipid mixing but failed to progress to pore formation [44]. In this system, fusion activity was restored when a hydrophilic cytoplasmic tail or a single arginine was added to the truncated HA construct, supporting a model in which the TMD must span the bilayer to promote fusion pore formation [44]. Similarly, fusion proteins in which the TMD was substituted by a glycosylphosphatidylinositol anchor are arrested in a hemifusion state, indicating that the progression to pore formation depends on perturbations of the inner membrane leaflets [45–47]. One proposed mechanism suggests that the TMD induces pore formation by generating spontaneous positive curvature in the hemifusion diaphragm [46], whereas another model proposes that TMD induces elastic stress within the bilayer or alters its orientation within the membrane, thereby promoting lipid rearrangements that drive pore formation [48]. Although it remains unclear why viruses may require distinct TMD length, the thickness of the viral membrane may contribute to these requirements, which in turn may either be directly modulated by the TMD, or may depend on the viral lipid composition [49]. Together, these findings support the notion that the adequate TMD length of ANDV Gc is critical for late stages of membrane fusion, particularly, the transition from hemifusion to fusion pore formation, consistent with reports on other viral fusion proteins.

Our results also demonstrate that specific residues within the ANDV TMD contribute to fusogenic activity. Substitution of the highly conserved residue S1121 with alanine restricted fusion to lipid mixing, whereas substitution of the less conserved S1126 only produced a reduction of fusion and pseudotyped vector cell entry. Residue-specific effects within TMD have also been reported for other viral fusion proteins. For example, a double mutation of glycine residues in the vesicular stomatitis virus glycoprotein resulted in a hemifusion phenotype, while the re-introduction of glycine at a different position restored fusion activity, suggesting a role in TMD-lipid interactions, rather than TMD-TMD contacts [50]. Glycine-dependent effects have been described for influenza HA, where mutation G520S was tolerated but G520L prevented fusion pore formation in strain A/Japan/305/57 (H2/N2) [51], whereas equivalent substitutions in strain X:31 (H3N2) were better tolerated [44], indicating strain-specific differences.

In the more-closely related fusion protein E1 of Semliki Forest virus, substitution of conserved glycine residues within the TMD did not impair fusion and assembly, but showed an increase in cholesterol requirement [19]. In a complementary study, replacement of glycine-containing segments in the E1 TMD reduced fusion activity and destabilized E1–E2 interactions, suggesting that TMD contacts may contribute to coordinated glycoprotein rearrangements during viral cell entry [18]. Similar roles for TMDs have been reported for flavivirus class II fusion protein E. The transmembrane hairpin is essential for membrane fusion and mutations within the TMD affect early fusion steps and the stability of the post-fusion E homotrimer [52]. TMD-mediated intra-trimer and inter-trimer interactions may also contribute to efficient membrane fusion and to protein rearrangements during viral maturation and particle assembly [53]. Residue-specific requirements within TMDs have additionally reported for other viral envelope proteins, including Moloney murine leukemia virus envelope glycoprotein, where proline substitutions reduced syncytia formation [54] and the HIV gp41, where replacement of a conserved arginine abolished syncytia formation [40]. In HIV gp41, this conserved arginine mediates heterotypic interactions with the TMIS, promoting lipid mixing and formation of the stable post-fusion homotrimer [55]. Similar TMIS-TMD interactions have been described for Ebola virus GP and parainfluenza virus 5 F class I proteins [56,57]. Together with structural studies of membrane-embedded TMDs [20,22,33,58], these observations indicate that TMDs of viral fusion proteins contain specific sequence determinants that regulate fusion protein stability and drive critical roles by orchestrating homotypic or heterotypic TMD-TMD or TMD-TMIS contacts.

Small residues such as glycine and serine are frequently found at helix–helix interfaces of TMDs, where they reduce the distance between helix axes [59–61]. Serine is particularly notable because its hydroxyl group can participate in hydrogen bonding by serving as donor or acceptor and it is the third most common disorder-promoting residue [62]. By incorporating an additional hydrogen bond, serine can introduce a bending angle in transmembrane helices, which can result in a significant displacement of residues located at the other side of the membrane, thereby transmitting conformational changes from the TMD to the ectodomain [63]. Although a functional role of serine residues has not been widely described for TMDs of other viral fusion proteins, our findings indicate that the conserved S1121 residue of ANDV Gc plays a critical role in late stages of membrane fusion. This effect may reflect conformational rearrangements within the fusion protein or intermolecular interactions required for efficient pore formation.

Interestingly, all fusion-inactive mutations also failed to support VLP assembly, further emphasizing a role of the Gc TMD in molecular interactions. Although the *in situ* structure shows contacts between the N-terminal region of the Gc TMD and Gn TMD2 [33], the length and specific residue requirement described here cannot be readily explained by this pre-fusion structure. Future structural studies addressing rearrangements during viral particle biogenesis and fusion intermediates may therefore provide further mechanistic insight.

## Acknowledgments

The authors used Chat GPT as text edition support. The authors have reviewed and edited the output and take full responsibility for the content of this publication.

---

**Fig S1.**
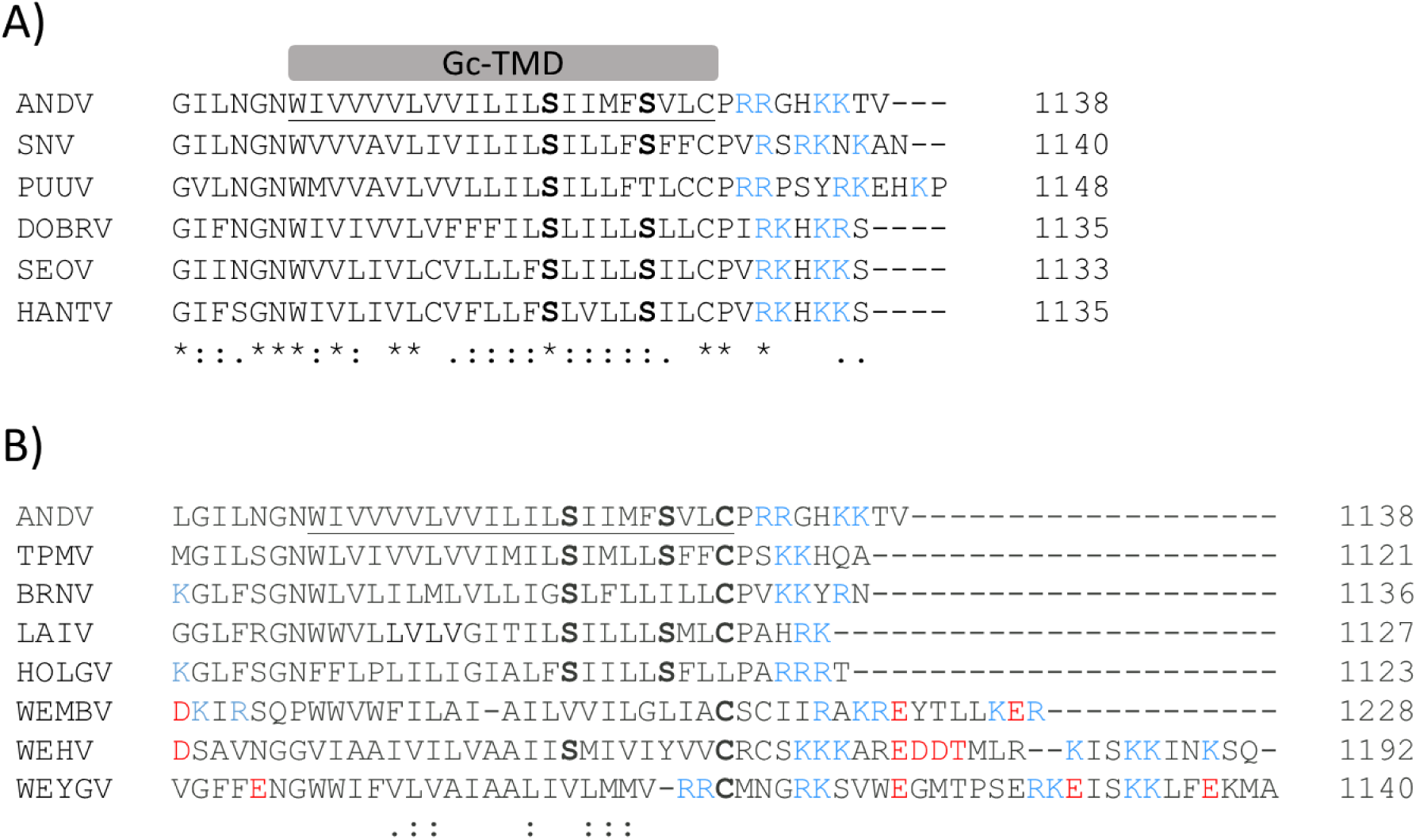
Multiple amino acid sequence alignment of the hantavirus glycoprotein precursor protein GPC from representative hantaviruses. Conserved residues are labeled below through point scheme; “:” indicates strongly conserved substitution while “.” indicates weakly conserved substitution. The predicted ANDV TMD sequence is underlined. Positively charged residues are indicated in blue, negatively charged residues in red. The conserved serine residues S1121 and S1126 (numeration from ANDV GPC) are indicated in bold letters. Numbers at the end of the alignment indicate the length of each respective GPC. **A)** Sequence alignment from GPC of representative pathogenic hantaviruses from the Orthohantavirus genus. ANDV (Andes virus, GenBank AAO86638.1), SNV (Sin Nombre virus, GenBank AAA75530), PUUV (Puumala virus, GenBank P21400, DOBRV (Dobrava virus NP_942554), SEOV (Seoul virus, GenBank NP_942557), HTNV (Hantaan virus, GenBank P08668. **B)** Sequence alignment from one representative protype of each of the eight genera of the Hantaviridae family. ANDV (Andes virus, Orthohantavirus genus (mammal hosts), GenBank: AAO86638.1), TPMV (Thottapalayam virus, Thottimvirus genus (mammal hosts), GenBank ABH09887.1), BRNV (Brno virus, Loanvirus genus (mammal hosts), GenBank APU53640.1), LAIV (Laibin virus, Mobatvirus genus (mammal hosts), Genbank AJZ68871.1), HOLGV (Hainan oriental leaf-toed gecko hantavirus, *Reptillovirus* genus (reptilian hosts), GenBank AVM87654.1), WEMBV (Wenling minipizza batfish hantavirus, *Actinovirus* genus (piscine hosts), GenBank: AVM87657.1), WEHV (Wenling hagfish virus, *Agnathovirus* genus, (piscine hosts), GenBank AVM87666.1), WEYGV (Wenling yellow goosefish hantavirus, *Perciloirus* genus (piscine hosts), GenBank AVM87660.1).

**Fig S2.**
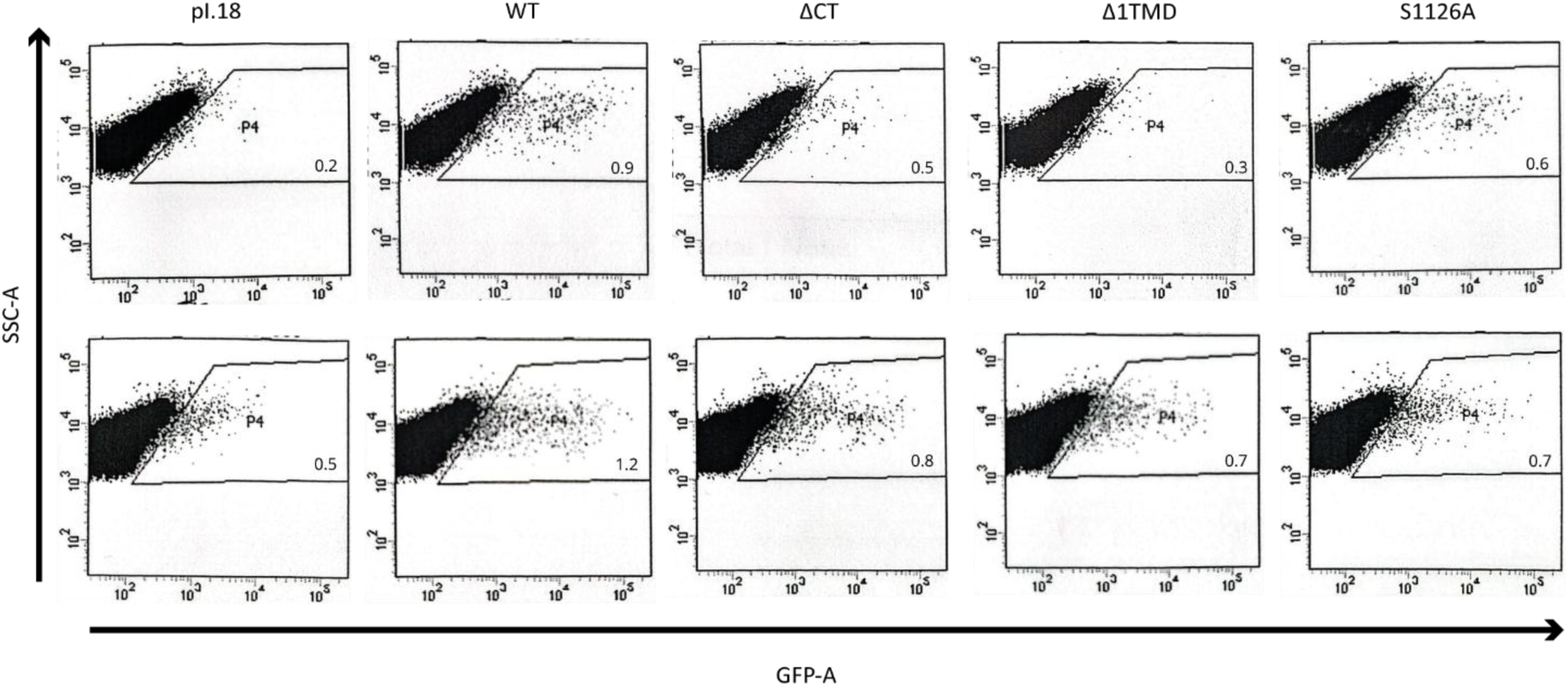
Raw data of cell cytometry dot blots of Vero E6 cells transduced with SIV vectors pseudotyped with ANDV WT or TMD mutants. SIV vectors pseudotyped with ANDV Gc WT or TMD were used to transduce Vero E6 and a GFP reporter gene expression measured 72 h post-transduction. Gates showing the percentage of cells expressing GFP were established using untransfected cells and cells transfected with the empty vector (pI.18) to rule out autofluorescence. Dot blots were acquired from N = 2 independent experiments. For the calculation of positive events, the negative control signal from pI.18 transfected cells was subtracted from the experimental samples to account for background fluorescence.

